# Melanocortin 4 Receptor-Dependent Mechanism of ACTH in Preventing Anxiety-Like Behaviors and Normalizing Astrocyte Proteins After Early Life Seizures

**DOI:** 10.1101/2024.10.09.617457

**Authors:** Mohamed R. Khalife, Colin Villarin, Juan Manuel Ruiz, Sam A. McClelland, Khalil Abed Rabbo, J. Matthew Mahoney, Rod C. Scott, Amanda E. Hernan

## Abstract

Epilepsy, affecting millions globally, often leads to significant cognitive and psychiatric comorbidities, particularly in children. Anxiety and depression are particularly prevalent, with roughly a quarter of pediatric epilepsy patients having a comorbid diagnosis. Current treatments inadequately address these issues. Adrenocorticotropic hormone (ACTH), a melanocortin peptide, has shown promise in mitigating cognitive deficits after early life seizures (ELS), potentially through mechanisms beyond its canonical action on melanocortin 2 receptor (MC2R). This study explores the hypothesis that recurrent ELS is associated with long-term anxiety, and that treatment with ACTH can prevent this anxiety through a mechanism that involves melanocortin 4 receptors (MC4R) in the brain. Our findings reveal that ACTH ameliorates anxiety-like behavior associated with ELS, without altering seizure parameters, in wildtype (WT) mice but not in MC4R knockout (KO) mice. Our findings also show that knocking-in MC4R in either neurons or astrocytes was able to rescue the anxiety-like behavior after ACTH treatment. Further, our results show that ACTH normalizes important astrocytic proteins like Glial Fibrillary Acidic Protein (GFAP) and Aquaporin-4 (AQP4) after ELS. This suggests that ACTH’s beneficial effects on anxiety are mediated through MC4R activation in both neuronal and astrocytic populations. This study underscores the therapeutic potential of targeting MC4R in epilepsy treatment, highlighting its role in mitigating cognitive impairments and anxiety-like behaviors associated with ELS.

## Introduction

Epilepsy, affecting 50-65 million people worldwide, is a neurological condition characterized by an enduring predisposition to generate seizures with significant neurobiological, cognitive, and social implications^1,2^. Approximately 30-40% of children with epilepsy suffer from cognitive, psychiatric or behavioral comorbidities, including a particularly high incidence of anxiety disorders^3–5^, which significantly impact quality of life^6–9^. Current anti-seizure therapies fail to adequately address these comorbidities^6,10,11^.

ACTH, a melanocortin family peptide within the hypothalamus-pituitary-adrenal (HPA) axis, has been utilized as a treatment for decades in childhood epilepsies, particularly infantile spasms^12–15^. The canonical mechanism of action of ACTH is through agonism of melanocortin 2 receptor (MC2R) in the adrenal glands, where it plays a role in steroidogenesis^16,17^. However, because ACTH agonizes all melanocortin receptors, receptors other than the canonical MC2R may be involved as well. Indeed, we showed that rodents with a history of early life seizures treated with ACTH, but not a corticosteroid, exhibited significant enhancements in fear extinction learning and attention^18,19^. We also showed that ACTH, but not a corticosteroid, normalized expression of genes in the brain involved in synaptic plasticity and cell communication and signaling, without changing seizure parameters^20^. Taken together, these data support a role for ACTH acting at an additional pathway, with actions above and beyond canonical corticosteroid release.

The melanocortin family of receptors, MCRs1-5, all bind ACTH with varying affinity and exhibit distinct tissue distributions and functions^18,21–24^. Activation of MC4R, primarily expressed in the CNS, has been shown to be neuroprotective in neurodegenerative diseases^25–27^. However, recent studies indicate that MC4R activation can decrease cell death, enhance cognitive performance in disease models, and regulate synaptic plasticity in the hippocampus^25–31^. Activating the hippocampal MC4R circuit alleviated synaptic plasticity impairments in an Alzheimer’s disease APP/PS1 double-transgenic mouse model^28^. Moreover, activation of MC4R enhances synaptic plasticity by regulating dendritic spine morphology and increasing the abundance of AMPA receptors^29^. This indicates that MC4R has therapeutic potential in conditions where synaptic function is compromised.

Research has mainly focused on synaptic and neuronal function after early life seizures (ELS) and it was shown that ELS can disrupt the hippocampal-prefrontal cortex network, alter synaptic plasticity and reduce neurogenesis, leading to cognitive and psychiatric deficits. However, there is a significant gap in the understanding of the role of astrocytes, which play a vital role in the brain’s response to injury, as well as maintaining ionic balance and neuronal signal transmission, after ELS. MC4R is also expressed in astrocytes. Activation of MC4R in astrocytes promotes anti-inflammatory effects and modulates cell survival proteins, highlighting its neuroprotective role^32^. Astrocytes are crucial in epilepsy, resetting ion balance and recycling neurotransmitters after seizure, and maintaining blood-brain barrier integrity^33^. Astrocyte channels, transporters, and metabolism alterations contribute significantly to seizure predisposition and epilepsy onset^34^. For instance, increased levels of Glial Fibrillary Acidic Protein (GFAP), indicative of astrogliosis, and altered Aquaporin-4 (AQP4) expression in astrocytes leading to disrupted water regulation, are both linked to increased seizure susceptibility and worsened seizure activity^33,35–39^. However, the role of MC4R activation in mitigating astrocyte dysfunction in epilepsy has not been explored.

We have previously shown in two rat models that ACTH can improve cognitive outcome after early life seizures^18,19^. Here we show for the first time that ELS is associated with significant anxiety in a mouse model, which is ameliorated by ACTH. In this study, we sought out to test whether ACTH was exerting its positive effects through MC4Rs in both neuronal and astrocyte populations in the brain. Our findings underscore a crucial role for MC4R in ACTH’s positive effects and its potential to alleviate anxiety-like behavior.

## Materials and Methods

### Animals

Male and female mice were used in all studies. C57BL/6J (strain#:000664), MC4r KO (B6.129S4-Mc4rtm1Lowl/J, Strain #:032518), Syn1-cre (B6.Cg-Tg(Syn1-cre)671Jxm/J, Strain #:003966) and Gfap-cre (B6.Cg-Tg(Gfap-cre)77.6Mvs/2J, Strain #:024098) mice from the Jackson Laboratory were used in the experiments. The mice were provided with food and water *ad libitum* and maintained on a 12-h light-dark cycle. All experiments were performed in accordance with the guidelines instated by the National Institutes of Health *Guide for the Care and Use of Animals*. The animal protocols were approved by the Institutional Animal Care and Use Committee at Nemours Children’s Health.

MC4R knockout mice were maintained as homozygous, and littermates with both copies of the receptor were obtained from Jackson Labs and used as controls. This strain has a floxed stop codon in front of the MC4R gene, allowing us to knock the receptor back using cre-recombinase. For the knock-in experiments, we used a breeding strategy with homozygous MC4R KO mice and hemizygous GFAP Cre or Syn1 Cre mice to re-express MC4R in astrocytes or neurons, respectively. Male hemizygous GFAP Cre, male hemizygous Syn1 Cre and homozygous KO female mice were used. We bred homozygous MC4R KO females with hemizygous GFAP Cre males in the first cross. We genotyped the F1 generation and selected heterozygous KO mice that were also Cre positive. The one floxed allele of the MC4R gene was reintroduced by the Cre recombinase, resulting in heterozygous MC4R knockout mice. F1 generation heterozygous KO/Cre+ mice were bred with homozygous MC4R KO mice. 75% of the offspring from breeding a heterozygous KO/Cre+ mouse with a homozygous KO mouse will be Cre positive. The Syn1 Cre mice were bred in a similar manner.

### Early Life Seizure Mouse Model

Seizures begin on postnatal day 10 (p10). 20 total seizures were administered for 5 days from p10-p14. Wild type and knockout mice were divided into three groups each: WT control vehicle-treated, vehicle-treated ELS, and ACTH-treated ELS; and KO control vehicle-treated, vehicle-treated ELS, and ACTH-treated ELS. (WT: N = 7 control vehicle-treated, N = 8 vehicle-treated ELS, N = 9 ACTH-treated ELS; KO: N = 7 control vehicle-treated, N = 7 vehicle-treated ELS, N = 6 ACTH-treated ELS). Knock-in mice were split into 2 groups each: (N = 6 MC4R Syn1 KI vehicle-treated ELS, N = 8 MC4R Syn1 KI ACTH-treated ELS, N = 5 MC4R GFAP KI vehicle-treated ELS, N = 5 MC4R GFAP KI ACTH-treated ELS) **(Table 1)**. An hour before every seizure, mice received subcutaneous injections of their respective drug or vehicle. Animals took a one-hour break between each of their four daily seizure sessions. 4-animal groups were housed in individual sections of a custom-designed and built plexiglass chamber, which were linked to a central chamber that contained flurothyl. All sections of the chamber have equal access to the central chamber containing the flurothyl when the section doors are closed, but a sliding plastic panel cuts off flurothyl access once the section is opened for quick evacuation of the flurothyl.

**Table 1.**
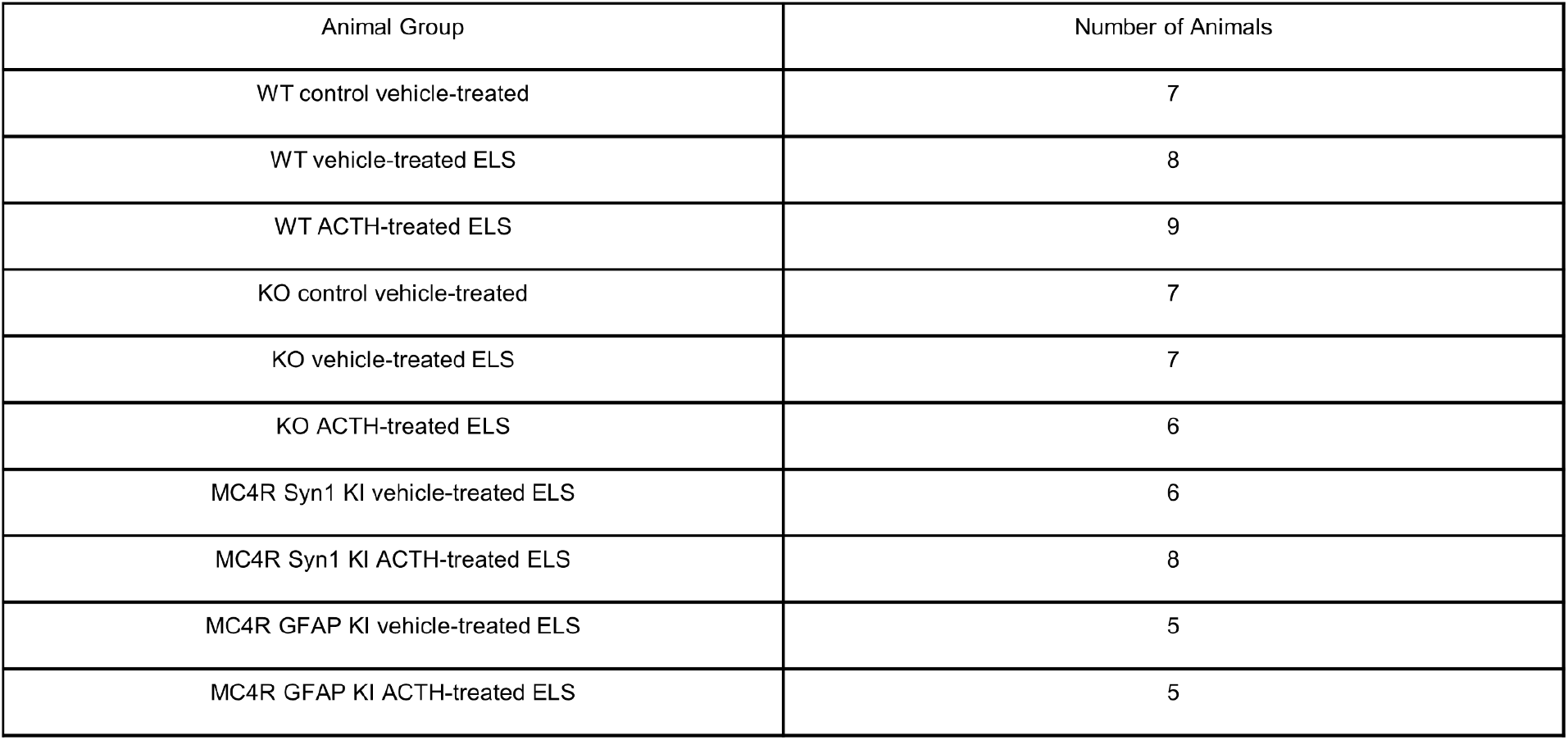
Wildtype, Knockout and Knock-in Mice Numbers.

| Animal Group | Number of Animals |
| --- | --- |
| WT control vehicle-treated | 7 |
| WT vehicle-treated ELS | 8 |
| WT ACTH-treated ELS | 9 |
| KO control vehicle-treated | 7 |
| KO vehicle-treated ELS | 7 |
| KO ACTH-treated ELS | 6 |
| MC4R Syn1 KI vehicle-treated ELS | 6 |
| MC4R Syn1 KI ACTH-treated ELS | 8 |
| MC4R GFAP KI vehicle-treated ELS | 5 |
| MC4R GFAP KI ACTH-treated ELS | 5 |

0.02mL of Flurothyl (Sigma-Aldrich) was dispensed into the central chamber on a filter paper and allowed to diffuse into the connected chambers containing individually placed animals. At incremental doses of 0.01mL or 0.02mL, flurothyl was given to elicit seizures with a minimum interval of one minute between each administration. The animal’s chamber was immediately evacuated upon the onset of a tonic-clonic seizure. Seizure sessions were recorded through webcams for close monitoring and further offline analysis. Animal-specific metrics, such as latency to seizure and duration of seizure, were documented by an observer and confirmed through video analysis.

### Drug Administration

Mice in the ACTH group received subcutaneous injections with an ACTH in 5% gelatin solution at a dose of 150 IU/m^2^, diluted with 5% gelatin to a total volume of 0.1mL. Vehicle Control and Vehicle ELS littermates received 0.1mL subcutaneous injections of the same solution vehicle as their littermates. All mice received drug administration once daily, one hour prior to each day’s seizure inductions.

### Open Field Task

At p50, mice were placed in a 46 cm x 46 cm squared arena in an isolated room and allowed to explore freely for 10 minutes. Sessions were video recorded and analyzed via ANY-maze for total locomotion, center entries as well as relative time spent in the center in order to obtain measures for spontaneous activity and exploration.

### Light/Dark Box Task

At p50, mice were placed in a 46cm x 27cm x 30cm acrylic box. 1/3 of the box was a dark area, while 2/3 of the box was exposed to light. A plexiglass door allowed the animals to enter either compartment of the box. The mouse was placed in the light compartment of the box and allowed to freely explore for 10 minutes. Animals were video recorded and analyzed via ANY-maze for time spent in each compartment in order to test anxiety-like behavior.

### Immunohistochemistry

48 hours after the last seizure, animals were perfused followed by brain collection. Brains were fixed with 4% paraformaldeyhde (PFA) for 4 hours and then transferred into 20% sucrose solution for cryopreservation. Brains were sliced at 40 microns for the prefrontal cortex (PFC) and hippocampus (HC) and fixed on charged slides for staining. A two-day immunohistochemistry protocol was used. Day 1 consisted of two 15-minute washes in 1X PBS, one hour of blocking and tissue permeabilization, followed by an overnight primary antibody incubation at 4C. Day 2 consisted of three 15-minute washes in 1X PBS, one two-hour secondary antibody incubation in the dark at room temperature, three additional 15-minute washes in 1X PBS, coverslip mounting using Fluoroshield with DAPI, and coverslip sealing. Blocking buffer was composed of 10% Normal Goat Serum and 0.5% Triton X-100 in 1X PBS. Tissues were co-stained with primary antibody, Glial Fibrillary Acidic Protein (GFAP) [1:1000] (ab4674, ABCAM LTD), and the conjugated secondary antibody, Alexa Fluor 555 (ab150170, ABCAM LTD), alongside another astrocytic biomarker Aquaporin-4 (AQP4) [1:500] (NBP1-87679, Novus Biologicals INC) and the secondary antibody, Alexa Fluor 488 [1:1000] (ab150077, ABCAM LTD). Stained sections were imaged at 40x using confocal microscopy, processing Z-stacks into maximum-intensity projections to visualize all tissue depths of fluorescent regions of interest (ROIs), and image analysis using ImageJ and the Colocalization Finder plug-in.

### Seizure Latency and Duration

To evaluate the differences in seizure latency and duration between various experimental groups, we conducted a time-to-event analysis using the Cox Proportional-Hazards model. We employed a shared frailty Cox Proportional-Hazards model due to its robustness in handling time-to-event data, allowing us to assess the effect of multiple covariates on the hazard of seizure occurrence adjusting for multiple measurements per animal. To account for day-specific effects, we stratified the Cox model by Seizure Day. This stratification ensures that our results are adjusted for any variations in seizure risk that may occur from day to day. This method enabled us to compare the seizure characteristics across different treatment and genotype groups, providing insights into the effects of various interventions on seizure dynamics.

### Statistical Analysis

The time spent in the center of the open field, center entries, total distance traveled and the time spent in the light compartment of the light dark box were analyzed using a generalized estimating equation (GEE) model, a multivariable, repeated-measures, regression model. This approach allows for modeling data with multiple measures per animal and adjusts for potential confounding variables like sex and litter. Sex, genotype and group were used as factors. Sex and litter were not significant and were therefore removed from the final analyses. Non-significance is defined as p>0.05; p values for all significant results are reported in the results section.

## Results

### Seizure Latency and Duration Is Unaffected by Treatment or MC4R knockout

We first asked whether our treatment differentially affected seizure induction in any of our groups. To do this, we analyzed seizure latency and duration. The mean latency and duration of seizures, along with their standard errors of the mean (SEM), were calculated for each group: WT ELS Vehicle (Latency: 344.73 ± 7.62 s, Duration: 68.06 ± 1.57 s), WT ELS ACTH (Latency: 298.52 ± 7.06 s, Duration: 72.97 ± 1.66 s), KO ELS Vehicle (Latency: 326.77 ± 9.59 s, Duration: 80.88 ± 3.47 s), KO ELS ACTH (Latency: 314.71 ± 9.69 s, Duration: 75.55 ± 2.31 s.) Cox Proportional-Hazards analysis was employed to assess the impact of treatment and genotype on seizure latency and duration, adjusting for multiple measurements per animal and day-specific effects. For seizure latency, the analysis revealed no significant effect of treatment (p = 0.73) or genotype (p = 0.81) across days **(Fig. 1A)**. For seizure duration, the analysis showed no significant effect of treatment (p = 0.16) or genotype (p = 0.22) across days **(Fig. 1B)**. Overall, our findings suggest that neither treatment nor genotype significantly influenced seizure latency or duration when adjusted for day-specific effects and multiple measurements per animal.

**Figure 1.**
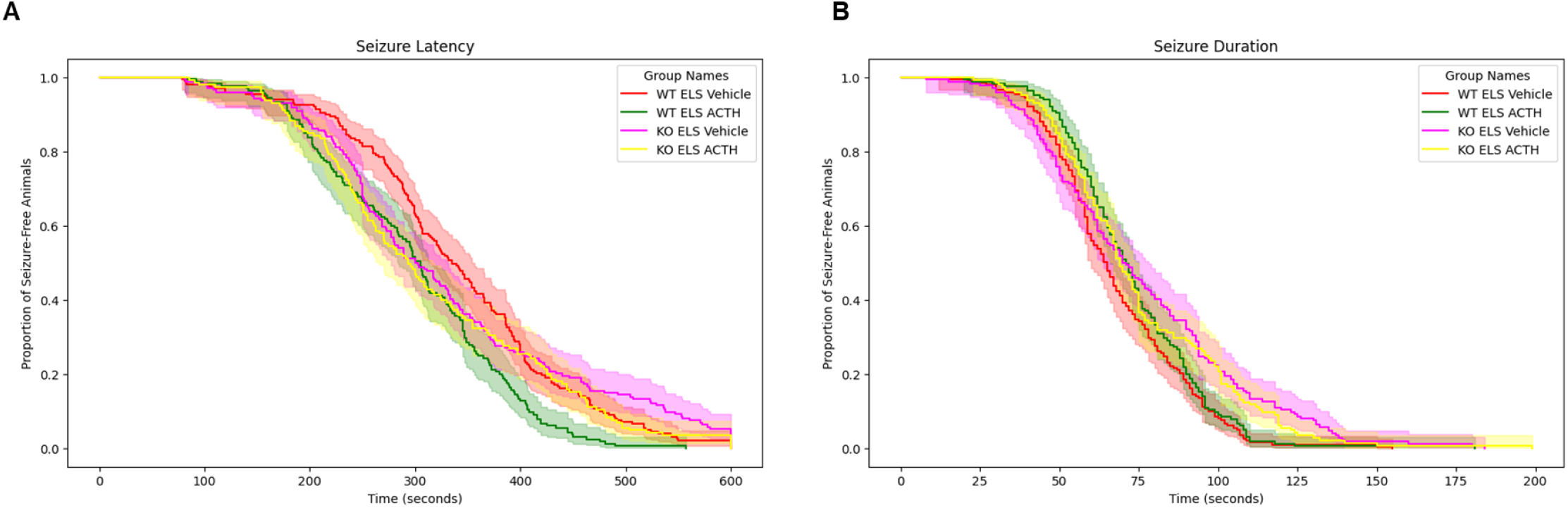
Seizure parameters are not altered by ACTH treatments. Treatment with ACTH does not alter latency to flurothyl seizure compared to vehicle-treatment groups across the different genotypes **(A)**. Similarly, treatment with ACTH did not significantly alter seizure duration across different genotypes **(B)**.

### ELS Did Not Affect Exploration and Spontaneous Activity in Open Field Task

The open field behavior test was conducted to assess spontaneous activity and exploration behavior in the experimental groups. This task measures the time spent in the center of the open field, the number of entries into the center, and the total distance traveled. These parameters provide insights into the animals’ willingness to explore a new environment and their activity. There were no significant differences in the time spent in the center of the open field between the control, vehicle-treated ELS, and ACTH-treated ELS (p > 0.05) (**Fig. 2A**). Similarly, the number of entries into the center did not differ significantly between the groups (p > 0.05) (**Fig. 2B**). Lastly, there was no significant variation in the total distance traveled by the animals across the different groups (p > 0.05) (**Fig. 2C**). These results indicate that neither treatment nor genotype had a significant effect on spontaneous activity or exploration in the open field task.

**Figure 2.**
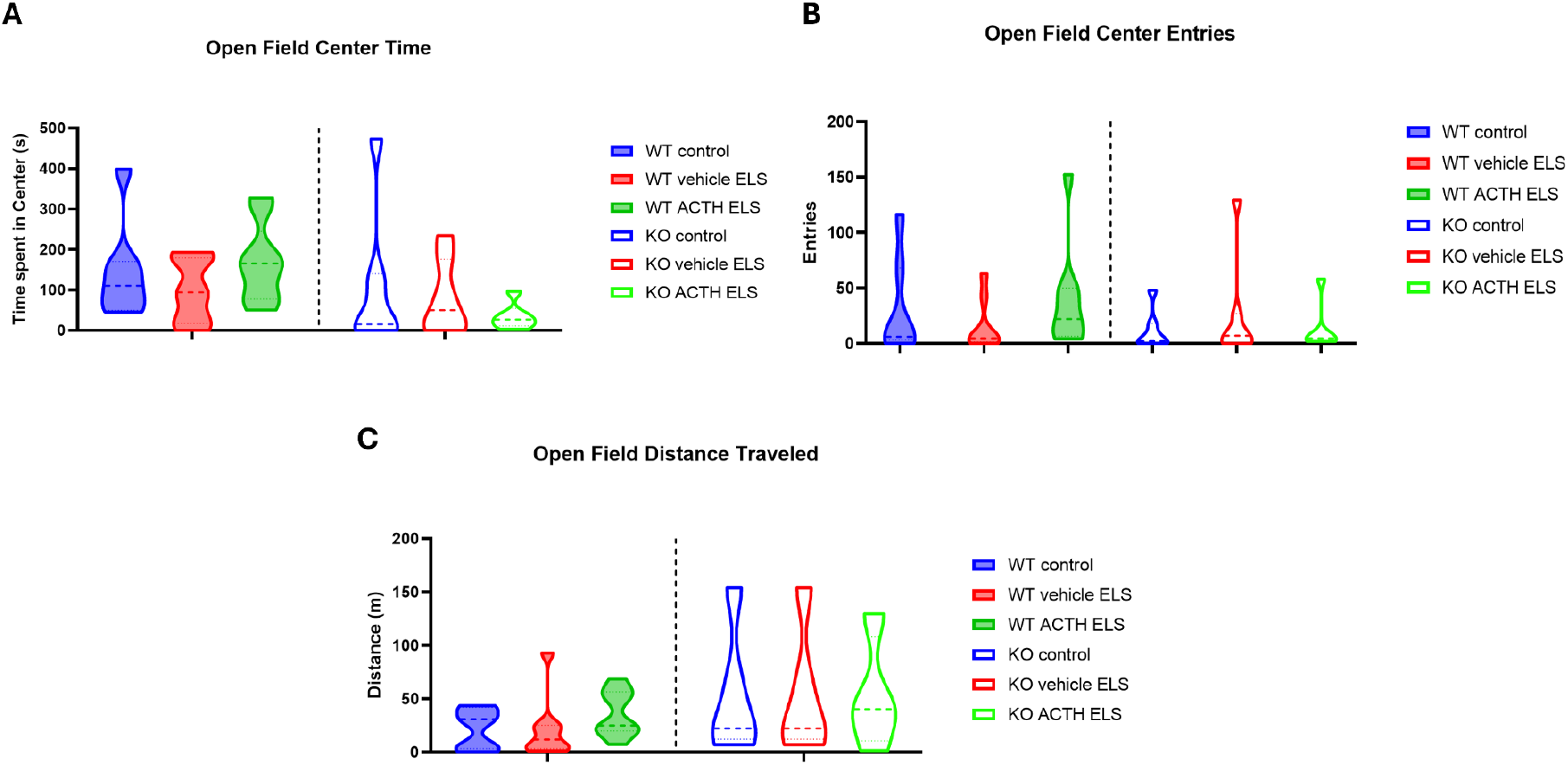
No Behavioral Differences in Open Field Task Parameters. There were no significant differences in the time spent in the center of the open field between the WT and KO groups **(A)**. Similarly, the number of entries into the center did not differ significantly between the WT and KO groups **(B)**. The total distance traveled by the subjects also showed no significant variation across the WT and KO groups **(C)**.

### ACTH Ameliorates Anxiety in Light/Dark Box Task

The light/dark box task was conducted to specifically assess anxiety-related behavior in the experimental groups. This task measures the time spent in the light zone of a box with both lighted and dark chambers, based on the natural aversion of rodents to brightly lit areas. Increased time in the light zone indicates reduced anxiety. Unlike the open field task, which primarily measures general locomotor activity and exploration, the light/dark box task is more sensitive to detecting anxiety-related behaviors.

WT vehicle-treated ELS mice spent significantly less time in the light zone compared to the WT controls (**Fig. 3;** p < 0.01). Interestingly, ACTH was able to ameliorate the anxiety in mice with a history of ELS.

**Figure 3.**
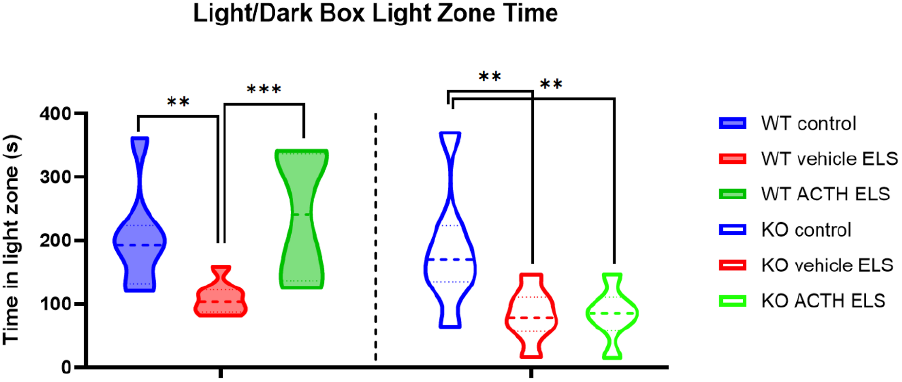
ACTH Ameliorates The Anxiety Phenotype In Mice With ELS. The light/dark box task was used to assess anxiety-related behavior by measuring the time spent in the light zone and the number of entries into the light zone. WT ELS mice treated with ACTH spent significantly more time in the light zone compared to WT ELS vehicle-treated mice. ACTH did not rescue the anxiety phenotype in MC4R KO ELS ACTH-treated mice compared to MC4R KO ELS vehicle-treated mice.

ACTH-treated ELS mice spent significantly more time in the light zone compared to the vehicle-treated group (p < 0.001). Similarly to the WT group, the MC4R KO vehicle-treated ELS spent significantly less time in the light zone compared to MC4R KO controls (p < 0.01), however, ACTH treatment was not able to ameliorate the anxiety (p < 0.01 to KO control group) indicating a MC4R-dependent mechanism for this amelioration, with a significant genotype*treatment effect (p<0.001).

### Cell-type specific re-expression of MC4R in neurons and astrocytes

MC4Rs are expressed on both neurons and astrocytes in the brain, where activation on each cell type can have differential effects on circuit function. We next asked whether MC4Rs on neurons or astrocytes were more important for the prevention of anxiety after ELS. To do this, we knocked-in (KI) MC4R into neurons or astrocytes using a syn1-cre or a GFAP-cre, followed by ELS and ACTH treatment. KI of the receptor did not change seizure parameters, and there were no significant effects of treatment on either the astrocyte- or neuron-re-expression group (**Fig. 4A,B** GFAP KI ELS Vehicle latency: 309.29 ± 9.28 s, duration: 79.02 ± 2.33 s; GFAP KI ELS ACTH Latency: 317.83 ± 10.86 s, Duration: 78.50 ± 2.64 s; Syn1 KI ELS Vehicle latency: 332.98 ± 8.89 s, duration: 82.04 ± 2.21 s; and Syn1 KI ELS ACTH latency: 347.32 ± 8.04 s, duration: 85.19 ± 2.14 s).

**Figure 4.**
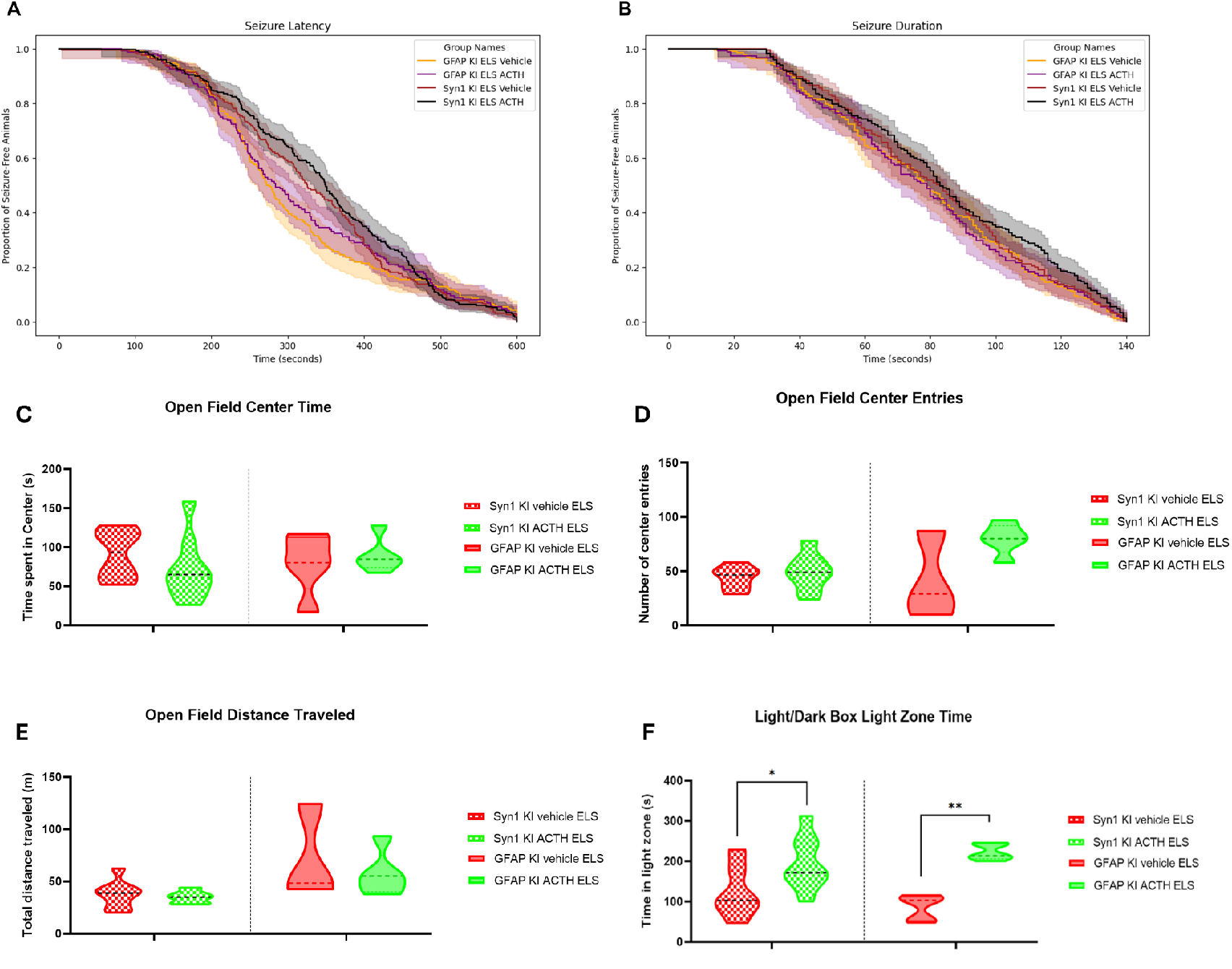
ACTH-Treated KI Mice With ELS Show Less Anxiety With No Differences in Seizure Parameters or Spontaneous Activity. Survival analysis plot indicates no differences in latency to tonic-clonic seizure between animals with MC4R re-expression in astrocytes and without treatment with ACTH (yellow vs purple line and envelope) or animals with MC4R re-expression in neurons with and without treatment with ACTH (red vs blue line) **(A)**. Similarly, treatment with ACTH did not significantly alter seizure duration in different groups of KI mice **(B)**. There were no significant differences in the time spent in the center of the open field between the vehicle treated and ACTH treated KI ELS groups **(C)**. Similarly, the number of entries into the center did not differ significantly between the vehicle treated and ACTH treated KI ELS groups **(D)**. The total distance traveled by the subjects also showed no significant variation across the vehicle treated and ACTH treated KI ELS groups **(E)**. Both KI ELS mice treated with ACTH spent significantly more time in the light zone compared to KI ELS vehicle-treated mice **(F)**.

We saw no significant differences in time spent in the center **(Fig. 4C)**, number of entries to the center **(Fig. 4D)** or the total distance traveled **(Fig. 4E)** between the vehicle-treated and ACTH-treated ELS knock-in (KI) groups in the open field. However, there is a genotype*treatment effect (p<0.001) for open field center entries, indicating that the treatment differentially affected the two reexpression groups. Post-hoc comparisons show that treatment with ACTH in animals where MC4Rs were re-expressed in astrocytes was significantly associated with an increase in the number of center entries compared to MC4R neuron KI groups (p<0.001 to both syn1-cre ELS vehicle and syn1-cre ELS ACTH groups) In the light/dark box task, ACTH treatment decreased anxiety in both the animals with MC4R re-expressed in neurons and animals with the receptor re-expressed in astrocytes **(Fig. 4F)**. Neuronal KI ELS mice treated with ACTH showed a significant increase in light zone time compared to neuronal KI mice treated with vehicle (p < 0.05). Similarly, the astrocyte KI ACTH-treated mice spent more time in the light zone compared to the vehicle-treated astrocyte KI ELS mice (p < 0.01). Notably, there is a genotype*treatment effect (p<0.001) for the light zone time. This was again driven by the ACTH-treated animals in the astrocyte-reexpression group, whose time spent in the light zone was significantly higher than all other groups (p<0.001). This suggests that re-expression in astrocytes was more effective at recovering the treatment effect than reexpression in neurons.

### Astrocyte dysfunction in the prefrontal cortex and hippocampus after recurrent early life seizures

Extensive research has focused on neuronal function, neuronal alterations and their long-term consequences following early life seizures. For instance, it was shown that early life seizures can disrupt the hippocampal-prefrontal cortex network and lead to cognitive deficits and psychiatric-like manifestations^40^. It was further shown that these deficits and manifestations occur due to changes in synaptic plasticity, such as an aberrant increase in long-term potentiation (LTP), rather than neuronal loss^40^. Other studies have also demonstrated that ELS does not result in cell death however it causes alterations in short- and long-term plasticity^41,42^ and reduced neurogenesis^43–47^. In addition to that, after ELS there is a decrease in inhibitory currents in hippocampus and neocortex, and hyperexcitation in neocortex^48,49^. While much is known about the neuronal alterations following ELS, there remains a significant gap in our understanding of astrocytic responses, their contributions to early life epilepsy and seizures, and their modulation of the MC4R effect. Given our data suggesting that re-expression of MC4R in astrocytes may be more effective than reexpression in neurons, we felt this gap was particularly important to address, as astrocytes are involved in various neuroprotective and neuroinflammatory processes that could influence seizure outcomes.

We used immunohistochemistry to examine astrocyte protein expression in the prefrontal cortex and hippocampus after ELS with and without ACTH treatment **(Fig. 5A)**. After seizures, we saw an increase in GFAP expression in both brain regions **(Fig. 5B,C)**, indicating enhanced astrocytic activation and gliosis. We also saw a decrease in AQP4 expression in both brain regions after ELS **(Fig. 5B,C)**. Dysregulation of AQP4 has been linked to increased seizure susceptibility through astrocyte proliferation, hypertrophy, impaired water balance, and edema ^33,35–39^. The results suggest that ELS is associated with astrocytic dysfunction. ACTH treatment was able to recover GFAP and AQP4 levels. These findings highlight the impact of ELS on astrocytic populations and the potential of ACTH treatment to mitigate these effects.

**Figure 5.**
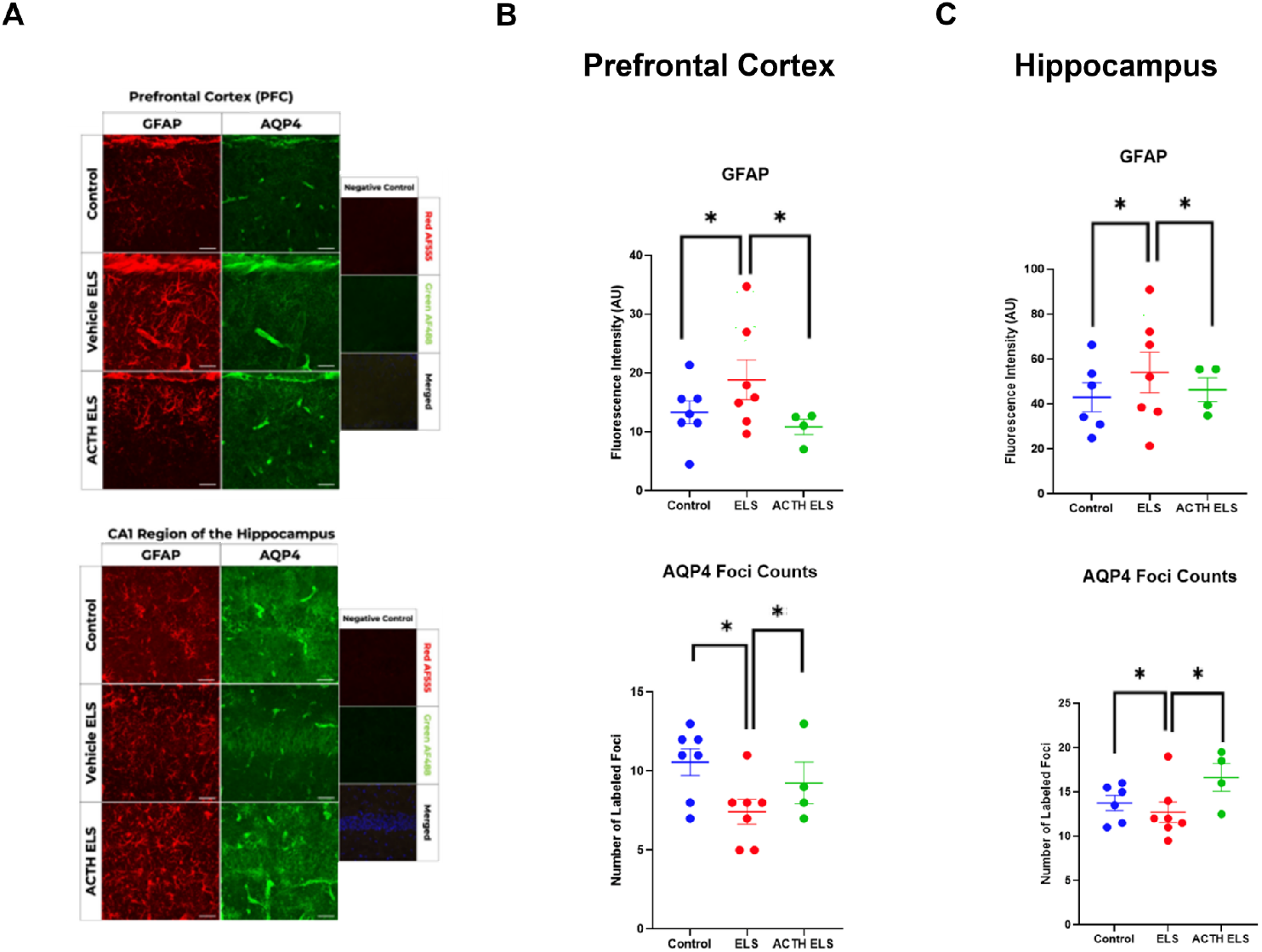
ACTH Mitigates The Astrocytic Dysfunction In Mice With ELS. Immunohistochemistry assessed astrocytic dysfunction after early life seizures **(A)**. Mice treated with ACTH showed a recovery in astrocytic function. ACTH normalized GFAP and AQP4 levels in the PFC after early life seizures **(B**,**C)**. ACTH normalized GFAP and AQP4 levels in the HC after early life seizures **(B**,**C)**.

## Discussion

The present study aimed to elucidate the mechanism by which ACTH can improve cognitive outcome after ELS. Cognitive deficits are a well-documented consequence of ELS, with up to 65% of patients experiencing impairments in anxiety, learning, memory, and executive function^6–9,50,51^. These deficits are particularly concerning as they can persist into adulthood, significantly impacting quality of life^6–9^. Traditional anti-seizure medications primarily focus on controlling seizure activity but do not address the cognitive comorbidities associated with epilepsy^6,10,11^. This gap in treatment highlights the need for therapeutic strategies that can mitigate the cognitive comorbidities. Our study investigated the effects of ACTH on anxiety, showing an MC4R-dependent improvement in anxiety with treatment, challenging the prevailing notion that the primary mediator of the effects of ACTH is through the MC2Rs in the adrenal cortex, leading to the systemic release of glucocorticoid steroid hormones^16,17^. In support of actions above and beyond corticosteroids, we previously showed that dexamethasone treatment failed to recapitulate the positive effects on behavior seen with ACTH treatment^18^ and that ACTH, but not dexamethasone, normalizes gene expression after ELS^20^.

ACTH is part of the melanocortin family of neuropeptide hormones, all of which arise from post-translational processing of proopiomelanocortin (POMC) and bind with varying affinity to the melanocortin receptors. MC2R is activated exclusively by ACTH, whereas the other 4 receptors can be activated by ACTH and all of the POMC neuropeptides, although the binding affinities differ. These G protein-coupled receptors also exhibit distinct tissue distributions and functions^18,21–24^. In the brain, melanocortin receptor MC4R was shown to have important roles in the pathophysiology of various neurological disorders like Alzheimer’s disease and cerebral ischemia. Activation of MC4R has been shown to promote neuroprotection and enhance cognitive function through multiple mechanisms. It was previously demonstrated that melanocortins protect against the progression of Alzheimer’s disease in triple-transgenic mice (3xTg); the study found that treatment with α-melanocyte-stimulating hormone (NDP-α-MSH) reduced cerebral cortex and hippocampus phosphorylation levels of amyloid/tau cascade proteins, inflammation, and apoptosis. Treated mice demonstrated decreased neuronal loss and improved learning and memory^31^. Another study, utilizing the TgCRND8 Alzheimer’s disease mouse model, found that α-MSH treatment prevents GABAergic neuronal loss and improves cognitive function. Treated mice exhibited improved spatial memory and reduced anxiety during the Y-maze task^52^. In the context of cerebral ischemia, MC4R agonists counteract late inflammatory and apoptotic responses post-ischemia in a transient global brain ischemia model. Treatment with MC4R agonists reduced the levels of pro-inflammatory cytokines and markers of apoptosis in the brain, leading to enhanced neuronal survival and functionality further improving cognitive performance in Morris Water Maze task^30^.

Furthermore, other studies demonstrated that activating MC4R has an impact on neuronal viability and synaptic plasticity. α-MSH treatment was shown to rescue neurons from excitotoxic cell death following kainic acid-induced damage resulting in a significantly higher number of viable neurons in the hippocampal CA1 pyramidal cell layer^53^. Moreover, activation of MC4R enhanced synaptic plasticity by increasing the number of mature dendritic spines and enhancing the surface expression of AMPA receptor subunit GluA1 mediating neurotransmission enhancement and hippocampal long-term potentiation^29^.

In our experiments, we utilized WT and MC4R KO mice. The use of the MC4R KO mice helps us understand the role of this CNS melanocortin receptor in mediating the positive effects of ACTH treatment on anxiety. Treatment with ACTH 1-hour before the start of each seizure day did not have significant influence on seizure latency or duration. This suggests that while ACTH may have therapeutic effects on other aspects of brain function, it does not appear to alter the fundamental characteristics of the seizures showing that ACTH is able to improve outcomes beyond altering the seizures^18,19^. This finding challenges the prevailing assumption that cognitive deficits are a result of the seizures.

The open field task results indicated that ELS did not significantly affect general locomotor activity or exploration behavior, as there were no differences in time spent in the center, number of entries into the center, or total distance traveled across the groups. This suggests that ELS and subsequent ACTH treatment do not impact the animals’ general activity levels or willingness to explore a new environment. In contrast, the light/dark box task revealed significant anxiety-related behaviors in ELS mice, which were ameliorated by ACTH treatment. WT ELS mice exhibited increased anxiety, spending less time in the light zone compared to controls. ACTH treatment significantly reduced this anxiety-like behavior, indicating its potential therapeutic effect. However, in MC4R KO mice, ACTH treatment did not ameliorate anxiety, showing that the anxiolytic effects of ACTH are mediated through MC4R signaling pathways.

The ability of ACTH to improve anxiety in WT mice but not in KO mice raised a question about the impact of neuronal and glial MC4R expression on these positive effects given that MC4R is expressed in both neurons and astrocytes. Astrocytes, glial cells in the brain, play a critical role in supporting neuronal function, modulating synaptic transmission and maintaining homeostasis within the central nervous system^33,54,55^. In epilepsy, astrocytes undergo significant alterations affecting astrocyte channels, transporters, and metabolism which are directly linked to epileptogenesis^34,56,57^. Specifically, disruptions in potassium, and water homeostasis, contribute to the seizure predisposition, epilepsy onset and the hyperexcitability characteristic of epilepsy^34,58,59^. Furthermore, astrocytes’ role in synchronizing neuronal activity and regulating synaptic transmission and plasticity, by releasing transmitters and maintaining calcium signaling, makes them essential for cognitive functions like learning and memory^54,55,60^. Disruptions in astrocytic function can lead to cognitive impairments^61^. Prolonged stimulation of hippocampal astrocytes, for instance, has been found to impair spatial memory, and working memory, indicating that astrocytic reactivity can negatively impact cognitive function^62,63^. However, sufficient activation of hippocampal astrocytes led to enhanced calcium activity sufficient to induce long-term potentiation (LTP) which is a cellular correlate of learning and memory. Mice performing contextual fear conditioning and T-maze tasks and undergoing hippocampal astrocyte activation during the acquisition phase demonstrated an improved memory recall in both tasks the following day^61^.

To address the question of cell-specific expression of MC4R, we re-expressed the MC4R selectively into neuronal populations or astrocytic populations. We observed in the open field task, the KI mice did not show significant differences in open field parameters. However, the significant genotype*treatment effect for the number of entries, with the GFAP KI*ACTH treatment showing a significant effect compared to the groups with MC4R neuron re-expression, highlights that the genotype influences the ACTH’s effect on exploration and activity levels in the open field. In the light/dark box task, we saw a decrease in the anxiety-like behavior in both the Syn1 KI and GFAP KI mice treated with ACTH. Notably, the significant genotype*treatment effect for the light zone time, with the GFAP KI*ACTH treatment showing a significant effect compared to all other groups, suggests that the effect of treatment on reducing anxiety was more pronounced in the GFAP KI genotype group shown by the increased time in the light zone for that group. This supports the role of MC4R in both neuronal and astrocytic populations in modulating anxiety responses following ELS. However, it is evident that the effects were more pronounced in the GFAP KI group. Therefore, the expression of MC4R in astrocytes appears to have a greater impact, with less anxiety, after ELS compared to its expression in neurons.

Immunohistochemistry staining was performed for important astrocytic proteins like GFAP and AQP4. The selected proteins play an essential role in astrocytic function and were shown to be altered in epilepsy. Increased levels of GFAP is observed in epilepsy and is indicative of astrogliosis^33,35^. AQP4 is a water channel protein involved in water homeostasis and balancing potassium concentration and its altered expression can lead to disrupted water regulation which is linked to increased worsened seizure activity^38,39^. This shows the importance of these proteins for proper astrocytic function. Our immunohistochemistry staining analysis demonstrated significant astrocytic dysfunction in both the prefrontal cortex and hippocampus following ELS, as evidenced by increased GFAP levels and decreased AQP4 expression. These changes indicate enhanced astrocytic activation and gliosis, as well as disrupted water channel function, which are consistent with previous research linking astrocytic alterations to increased seizure susceptibility and impaired neural function. ACTH treatment effectively normalized GFAP and AQP4 levels, suggesting its potential to mitigate astrocytic dysfunction induced by ELS. The recovery of these astrocytic markers highlights the therapeutic potential of ACTH in restoring normal astrocytic function and, by extension, improving neural homeostasis.

The findings of this study underscore the complex interplay between ELS, astrocytic function, and behavioral outcomes. The differential effects of ACTH treatment on seizure characteristics, anxiety-related behaviors, and astrocytic markers suggest that while ACTH may not directly influence seizure parameters, it holds promise in ameliorating anxiety and astrocytic dysfunction associated with ELS. Future research should focus on elucidating the precise mechanisms through which ACTH exerts its therapeutic effects.

